# Cortex-wide spatiotemporal motifs of theta oscillations are coupled to freely moving behavior

**DOI:** 10.1101/2024.09.27.615537

**Authors:** Nicholas J Sattler, Michael Wehr

**Affiliations:** University of Oregon

**Author notes:** Corresponding Author: Michael Wehr, Institute of Neuroscience, 1254 University of Oregon, Eugene, OR 97403.

## Abstract

Multisensory information is combined across the cortex and assimilated into the continuous production of ongoing behavior. In the hippocampus, theta oscillations (4-12 Hz) radiate as large-scale traveling waves, and serve as a scaffold for neuronal ensembles of multisensory information involved in memory and movement-related processing. An extension of such an encoding framework across the neocortex could similarly serve to bind disparate multisensory signals into ongoing, coherent, phase-coded processes. Whether the neocortex exhibits unique large-scale traveling waves distinct from that of the hippocampus however, remains unknown. Here, using cortex-wide electrocorticography in freely moving mice, we find that theta oscillations are organized into bilaterally-symmetric spatiotemporal “modes” that span virtually the entire neocortex. The dominant mode (Mode 1) is a divergent traveling wave that originates from retrosplenial cortex and whose amplitude correlates with mouse speed. Secondary modes are asynchronous spiral waves centered over primary somatosensory cortex (Modes 2 & 3), which become prominent during rapid drops in amplitude and synchrony (null spikes) and which underlie a phase reset of Mode 1. These structured cortex-wide traveling waves may provide a scaffold for large-scale phase-coding that allows the binding of multisensory information across all the regions of the cortex.

**Bulleted list of key results:**

- Cortical theta oscillations are organized into bilaterally-symmetric spatiotemporal modes that span the neocortex.
- The dominant mode appears as a divergent traveling wave that originates in retrosplenial cortex and is correlated with mouse speed.
- Secondary modes are asynchronous spiral waves centered over somatosensory cortex.
- Secondary modes become prominent during transient drops in synchrony that underlie a phase reset of the dominant mode.
- We hypothesize that spiral waves may provide a mechanism to exert large-scale phase separability, and assimilate information into ongoing multisensory processing across the neocortex.

## Introduction

How are neural activity and computation coordinated across the brain, and how do they underlie behavior? Recent studies have highlighted the existence of prominent movement-related signals throughout the brain (Musall et al., 2019; Parker et al., 2020; Schneider and Mooney, 2018; Stringer et al., 2019). This widespread activity likely reflects the integration of sensory processing with ongoing movement, which is dynamically modulated as animals interact with and actively sample the sensory environment during natural freely moving behavior (Ferezou et al., 2006; Grion et al., 2016). In the broader context of brain function, processes such as sensorimotor integration, decision-making, and memory rely on the ability of neural circuits to communicate flexibly and efficiently across widespread brain regions. Although local and long-range connections undoubtedly provide the anatomical substrate for this communication, it is less clear whether fixed anatomical connections alone can meet the demands of dynamic and flexible communication required for the rapid timescale of natural behavior. Coherent oscillations have been proposed as a possible mechanism for such flexible routing of information across brain regions (Fries, 2005).

Oscillatory brain activity occurs across a wide range of frequencies, but the hippocampal theta rhythm (4-12 Hz) is especially prominent, and has been linked to locomotion and exploratory behavior in many species from rodents to humans (Buzsáki, 2002; M Aghajan et al., 2017). In particular, hippocampal theta plays a key role in spatial navigation, and memory formation, and coordinates neuronal spiking in other brain regions such as frontal cortex (Benchenane et al., 2010; Hyman et al., 2005; Jones and Wilson, 2005a, 2005b; Kim et al., 2011; Siapas et al., 2005). The theta rhythm is not synchronous across the hippocampus, but rather forms a traveling wave along the septotemporal axis (Lubenov and Siapas, 2009; Patel et al., 2012; Zhang and Jacobs, 2015). Traveling theta waves have also been observed in human intracranial cortical recordings, where they are related to memory encoding and recall (Bahramisharif et al., 2013; Mohan et al., 2024; Zhang et al., 2018). While theta and respiratory-entrained oscillations have been recorded across the neocortex (Karalis and Sirota, 2022; Tort et al., 2018), whether they are spatiotemporally organized into traveling waves, or how they are related to behavior, remains unknown.

Recent studies have demonstrated that cortical activity exhibits metastable dynamics, in which brain activity unfolds as a sequence of discrete quasi-stationary states that are separated by abrupt transitions (La Camera et al., 2019; Recanatesi et al., 2022). For example, spatiotemporal patterns of gamma oscillations across the cortex appear to occur in bouts, which contain stimulus information and are separated by brief transitional periods of low-amplitude, asynchronous activity that have been termed “null spikes” (Freeman, 2007; Kozma and Freeman, 2017). Metastable state sequences of cortical spiking patterns conveying stimulus information have similarly been observed across species, and have been theorized to allow neural computations to proceed along a sequence of distinct epochs, in which brain activity forms quasi-stable neural representations punctuated by rapid state transitions (Abeles et al., 1995; Jones et al., 2007; Ponce-Alvarez et al., 2012). With a lack of direct recordings, whether metastable dynamics occur for traveling waves of theta oscillations, or what the spatial extent of the metastable states may be, has remained theoretical (Cabral et al., 2022; Roberts et al., 2019).

To investigate these spatiotemporal dynamics at a cortex-wide scale, we recorded field potentials using a large-scale electrocorticography (ECoG) array covering nearly the entire dorsolateral surface of the neocortex in freely moving mice. We found prominent theta oscillations that were spatiotemporally organized into three major bilateral modes that were strongly modulated by mouse running speed. These three patterns of activity showed metastable dynamics, alternating during transient desynchronization events which induced a phase reset in the dominant traveling wave pattern. These distinct cortex-wide spatiotemporal patterns of oscillatory activity could serve to integrate sensory, cognitive, and behavioral information that is distributed across the cerebral cortex.

## Results

### Theta oscillations are prominent across the cortex

We recorded local field potentials (LFPs) across the dorsolateral surface of the cortex using a 31-channel ECoG array (Figure 1A-C) while mice (N = 3) freely explored an open arena (Figure 1D). To monitor fine movements of the ears, nose, and face under freely moving conditions, we used an array of 3 head-mounted cameras (Figure 1D; (Sattler and Wehr, 2020)). LFP signals recorded from all channels and cortical regions exhibited a distinct peak in the theta range (Figure 1E,F). These theta oscillations showed pronounced phase differences across the cortical surface, consistent with a traveling wave. The instantaneous spatial phase gradient of theta-filtered LFPs (Figure 1G, time point corresponding to the vertical line in Figure 1E) typically appeared as a divergent traveling wave, indicating that moment-to-moment theta oscillations show cortex-wide spatiotemporal structure.

**Figure 1:**
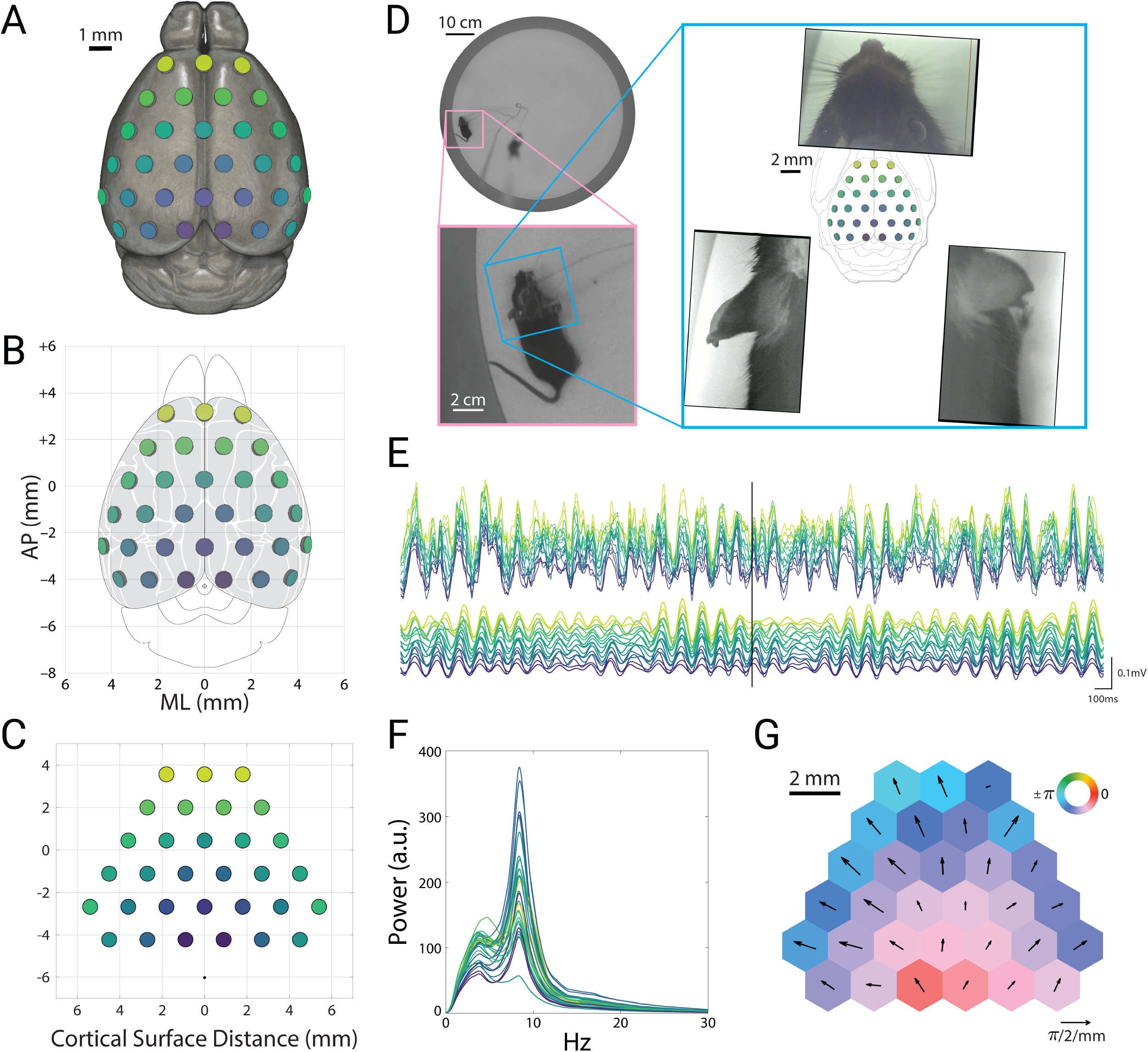
Theta oscillations are prominent across the cortex. A) Illustration of electrode placements of the electrocorticography (ECoG) array on the mouse brain (mesh adapted from alleninstitute.org). Colors indicate electrode position and correspond to channel colors in E-F. B) Electrode locations in conventional stereotactic coordinates, overlaid on the cortical parcellation of mouse brain atlas adapted from the Paxinos atlas (Kirkcaldie et al., 2012). C) Geometry of electrode array, showing electrode size (0.8mm) and inter-electrode distance (1.8mm). Black dot indicates the reference electrode location. Coordinates indicate inter-electrode distances along the curved surface of the skull, relative to initial alignment with bregma. D) Top view of the arena (top left), with inset showing an enlarged view of the mouse (bottom left). Overhead view of the head and ears from head-mounted cameras (right), with the ECoG array overlaid over the Paxinos atlas in the center. E) Example recording traces of the local field potentials (top) and theta-filtered signals (9 Hz, bottom). Vertical black bar corresponds to the time point shown in panels D and G. F) Power spectral density for each channel. Signals were band-pass filtered 2.5-30 Hz. G) Instantaneous phase (colors) and spatial phase gradient (quivers) of the theta-filtered signals at the time point indicated by the black bar in panel E. Hexagonal tiles indicate relative electrode positions, enlarged and reshaped for ease of visualization.

This instantaneous snapshot of a cortex-wide traveling wave (Figure 1E) closely matched the long-term average of the signals. The spatial phase gradient of these oscillations averaged across sessions and mice appeared as a bilaterally divergent vector field that originated in posterior retrosplenial cortex (Figure 2A, top). The mean amplitudes also showed a bilaterally symmetric pattern, with the highest theta amplitude in anterior retrosplenial cortex, roughly above the hippocampus (Figure 2A, bottom). Thus both the instantaneous and long-term average of these oscillations appeared as a global traveling wave. We wondered whether this traveling wave was stable over time, or showed dynamics that might relate to ongoing behavior. To address this, we turned to spectral decomposition methods to characterize spatiotemporal activity patterns.

**Figure 2:**
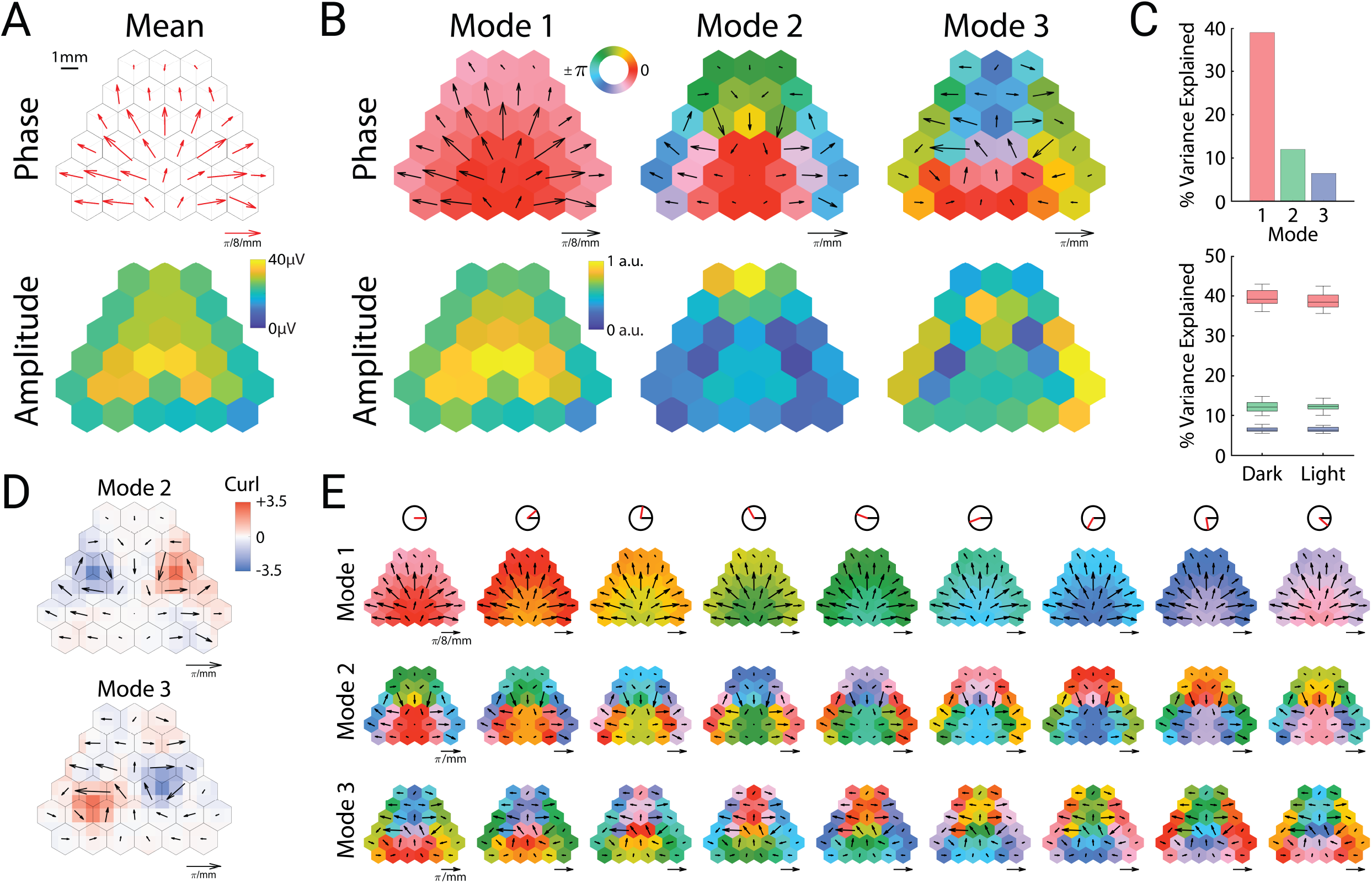
Theta oscillations can be decomposed into cortical modes. A) Mean spatial phase gradient (top) and amplitude (bottom). B) Phase (top row) and amplitude (bottom row) of the three prominent cortical modes. C) Variance explained by each mode across all sessions (top) and between dark and light sessions (bottom). Upper and lower portions of the box plots correspond to the upper and lower quartiles respectively. The middle lines correspond to the median value, and error bars correspond to minimum and maximum values. D) Vector curl calculated on interpolated values of the spatial phase gradients for modes two and three. E) Cortical cyclograms showing the progression of phase along their spatial phase gradients across one full cycle (from left to right). The red line in the top row indicates the current phase angle of the column.

### Theta oscillations can be decomposed into cortical modes

To identify distinct spatiotemporal motifs of oscillatory activity, we used complex-valued singular value decomposition to extract oscillatory “modes” (eigenvectors), that is, specific spatiotemporal patterns consisting of fixed amplitude and phase relations among the channels. This analysis revealed three prominent modes (Figure 2B), which collectively represented roughly 50% of the variance in the dataset, and were consistent in power across both illuminated and dark arena conditions (Figure 2C). These three cortical modes were strongly consistent across individual mice (Extended Data Figure 2-1).

The most prominent mode (“Mode 1”) strongly resembled the mean amplitude and phase gradient, and represented around 40% of the variance (Figure 2C). This oscillatory mode was not synchronous across the cortex, but rather showed a smoothly varying pattern of phase lags that formed a bilaterally divergent traveling wave originating near posterior retrosplenial cortex. In contrast to this pattern, the following two modes (Modes 2 and 3) were fully asynchronous (containing phase angles across the entire unit circle), with spatial phase gradients that formed bilateral spirals centered roughly over somatosensory cortex (Figure 2B).

**Extended Data Figure 1:**
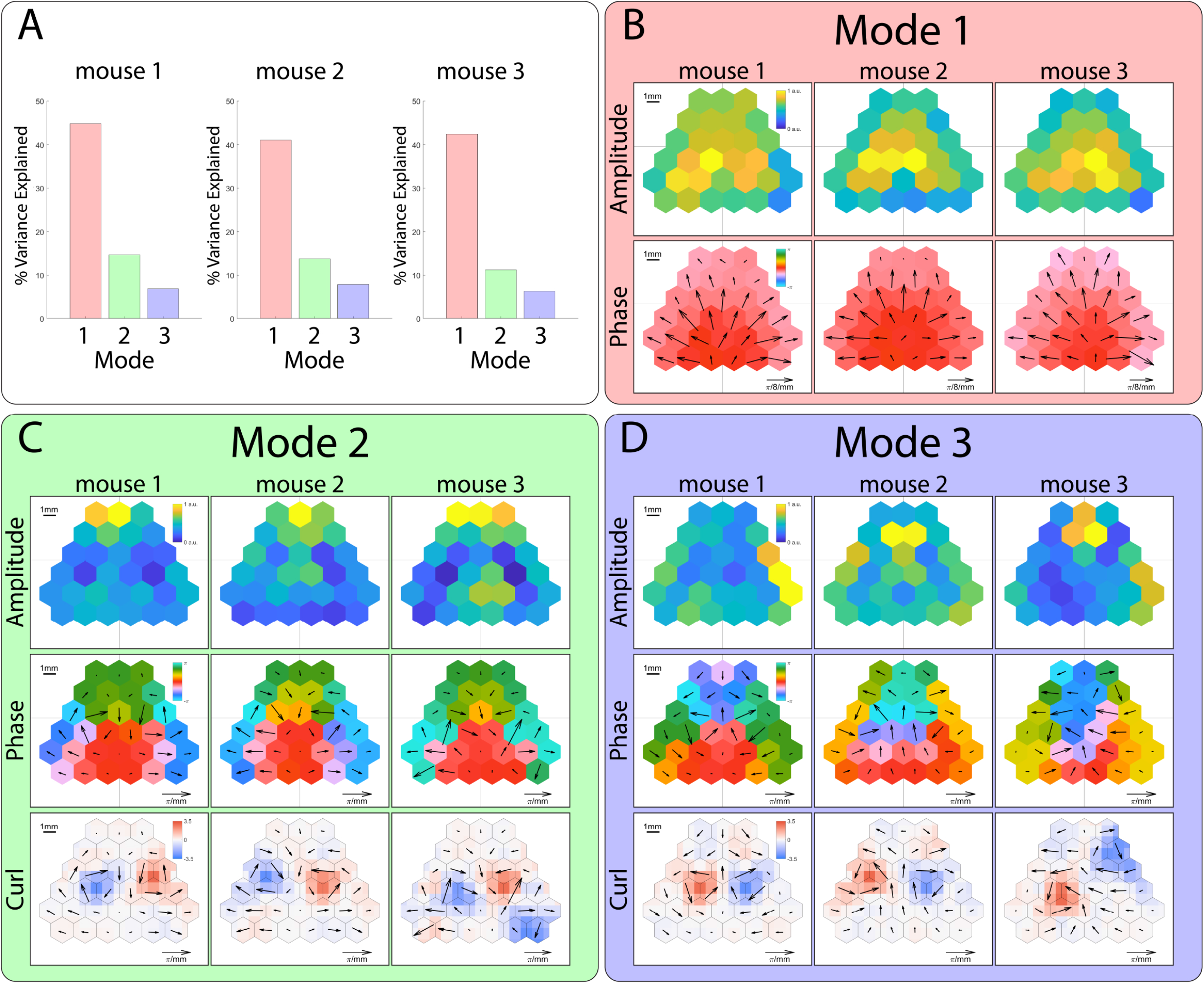
Cortical theta modes are robust across mice. A) Variance explained by each mode across all sessions for each mouse, quantified through singular value decomposition computed individually on each mouse. B) Amplitude (top row) and phase (bottom row) of Mode 1 for each mouse. C) Amplitude (top row), phase (middle row), and vector curl (bottom row) of Mode 2 for each mouse. D) Amplitude (top row), phase (middle row), and vector curl (bottom row) of Mode 3 for each mouse.

To quantify these spiral patterns, we computed the vector curl of Modes 2 and 3, which revealed strong bilateral singularities of opposite rotational directions (Figure 2D). Modes 2 and 3 differed in that Mode 2 was clockwise in the left hemisphere, whereas Mode 3 was counterclockwise (and vice versa). Similar to smaller mesoscopic spiral waves that have been previously observed in rodent (Huang et al., 2010, 2004) and human (Xu et al., 2023) neocortex, the lowest amplitudes were observed near the phase singularity for both modes. The location of the spiral waves, centered over somatosensory cortex, also matches the location of recently reported large-scale spiral waves in the 2-8 Hz frequency band in mice (Ye et al., 2023). Mode 2 displayed high amplitude in frontal cortex, whereas Mode 3 displayed high amplitudes in the temporal and parietal poles of the cortex (Figure 2B).

The bilateral phase symmetries of both Modes 2 and 3 result in opposite rotational directions of the left and right spirals. The temporal sequence of the phase gradient can therefore be described relative to the midline for both hemispheres. In the case of Mode 2, the phase sequence flows in a lateral → anterior → medial → posterior sequence, whereas Mode 3 flows in an anterior → lateral → posterior → medial sequence. Cortical cyclograms illustrate how the phase sequence of the modes unfold in time through a full theta cycle (Figure 2E and Supplementary Video 1).

### Modes are interrelated and correlate with mouse speed

The spatiotemporal motifs shown in Figure 2 represent time-averages across all recording sessions and mice. How do these modes unfold in time? To investigate the dynamics of these modes, we computed the instantaneous mean amplitude across all channels as a measure of overall cortical power, and the order parameter as a measure of synchrony across channels. The order parameter (see Methods) is a vector strength measure that varies from 0 (complete asynchrony) to 1 (complete synchrony).

A comparison of mean amplitude and order parameter over a representative ten-second period (Figure 3A top) along with the three modes (Figure 3A bottom) reveals strong relationships between these measures. Mean amplitude typically remained high for periods of roughly 1 second, and these periods were punctuated by brief drops in amplitude. These amplitude dips were tightly associated with sharp drops in the order parameter, indicating that global theta oscillations alternate between stable periods of high amplitude and synchrony, interrupted by brief events during which the amplitude and synchrony dropped to very low values. Similar brief desynchronization events have been previously described in ECoG recordings of gamma-band activity from local brain regions such as visual cortex or olfactory bulb, for which Freeman coined the term “null spikes” (Freeman, 2007; Kozma and Freeman, 2008). Although mean amplitude and order parameter showed distinct distributions (Figure 3B), cross-covariation analysis revealed that amplitude and order parameter were highly correlated with each other (Figure 3C, r = 0.605; p = 10^−9^, n = 48 trials), as expected from the large and tightly coordinated impact of null spikes on both measures. Moreover, Mode 1 was also strongly correlated with mean amplitude (r = 0.825; p = 10^−9^), whereas Modes 2 and 3 were anti-correlated with these measures (Figure 3C, Mode 2: r = -0.305; p = 10^−9^, Mode 3: r = -0.277; p = 10^−9^).

**Figure 3:**
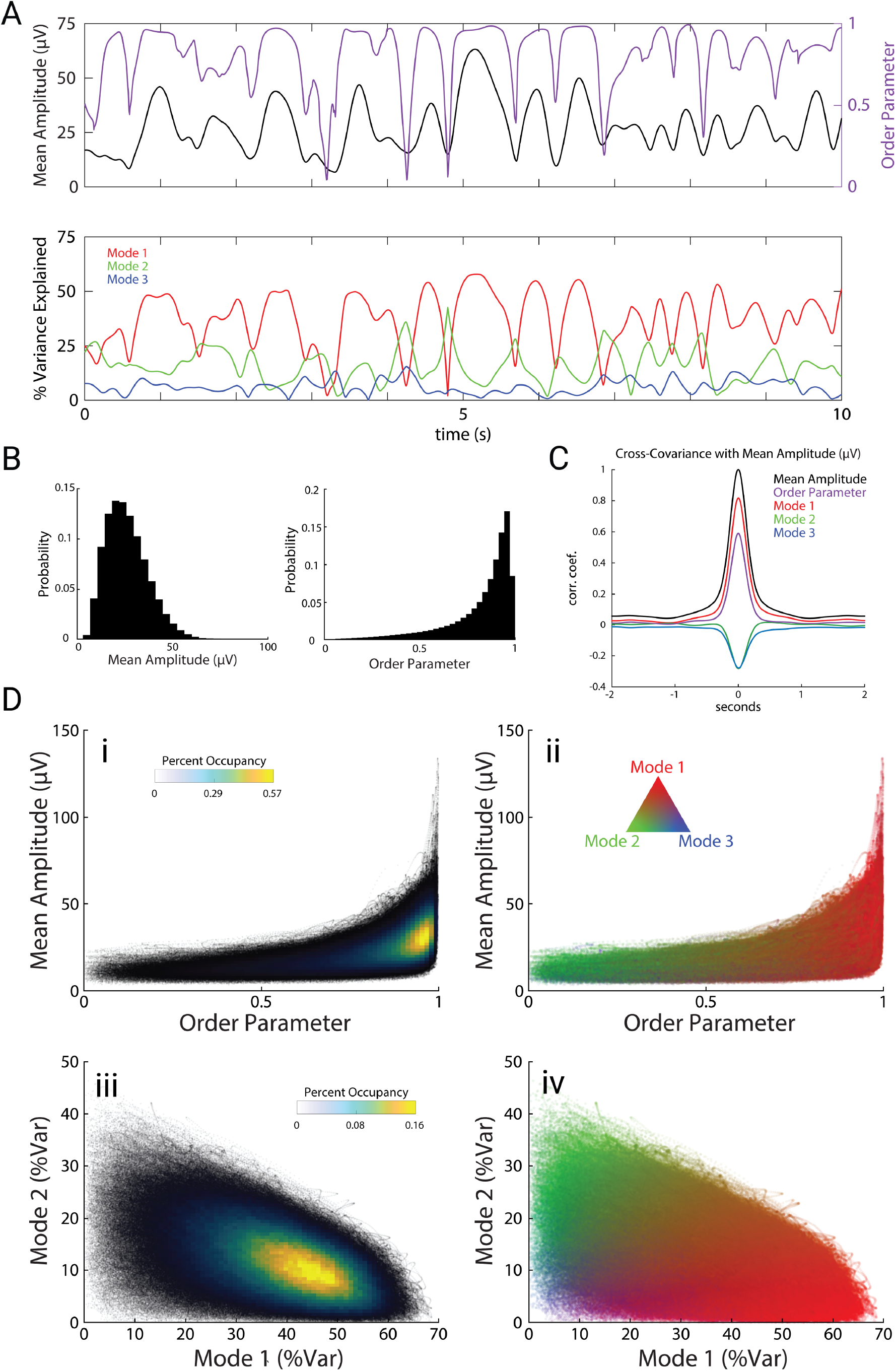
Modes are interrelated and correlate with amplitude and synchrony. A) Example trace of the mean electrode amplitude across channels (black), order parameter (purple), and modes 1, 2, and 3 (red, green, and blue respectively) across a 10 second period. Colors carry over across the figure. B) Distributions of mean amplitude and order parameter. C) Cross-covariance of the order parameter and modes (percent variance explained) with mean amplitude. D) Scatter plots with overlaid occupancy values (left) and their corresponding modal compositions (right).

The joint probability distribution of order parameter and mean amplitude shows that theta oscillations dwelled for most of the time at moderate levels of amplitude and synchrony (Figure 3Di), in which Mode 1 dominates (Figure 3Dii). Brief periods when theta activity plunged into low-order and low-amplitude null spikes (Figure 3Di) were dominated by the spiral pattern of Mode 2 (Figure 3Dii). However, Mode 2 activity was not confined to null spikes, but was consistently present during stable periods as well, coexisting in superposition with Mode 1 (Figure 3Dii, iii, iv).

These measures of theta oscillations were tightly correlated with mouse speed during free exploration of the open arena (Figure 4A left). Specifically, mouse speed was correlated with mean amplitude (r = 0.165; p = 10^−9^), order parameter (r = 0.067; p = 10^−7^), and Mode 1 (r = 0.126; p = 10^−9^). This indicates that these measures of global brain activity are related to mouse movements. In contrast to experiments with stereotyped mouse trajectories on linear tracks, open-field exploration involves wide variations in mouse speed, mouse turning, and coordinated movements of the head, ears, and nose. To investigate how these aspects of mouse movement related to theta oscillations, we first measured mouse turning, which we quantified as angular velocity ω (rads/s) of the mouse’s head. Mouse turning during open exploration was tightly correlated to speed (Figure 4A, r = 0.137; p = 10^−9^). During these turning events, the movements of the nose (left nostril: r = 0.130; p = 10^−9^, right nostril: r = 0.143; p = 10^−9^) and the ears (left ear: r = -0.0498; p = 10^−8^, right ear: r = -0.0684; p = 10^−9^) were tightly correlated with the direction of the turn (Figure 4B). Whereas turning was correlated with speed, and speed was correlated with theta activity (amplitude, order parameter, and Mode 1), we found that turning was not correlated with any measures of theta (Figure 4A right, r = 0.008, 0.0009, -0.005; p = 0.069, 0.789, 0.238). Ear and nostril movements were also uncorrelated with any measures of theta (data not shown). Theta activity therefore appeared to be specifically related to locomotion speed as opposed to general movements of the mouse, head, ears, or nose.

**Figure 4:**
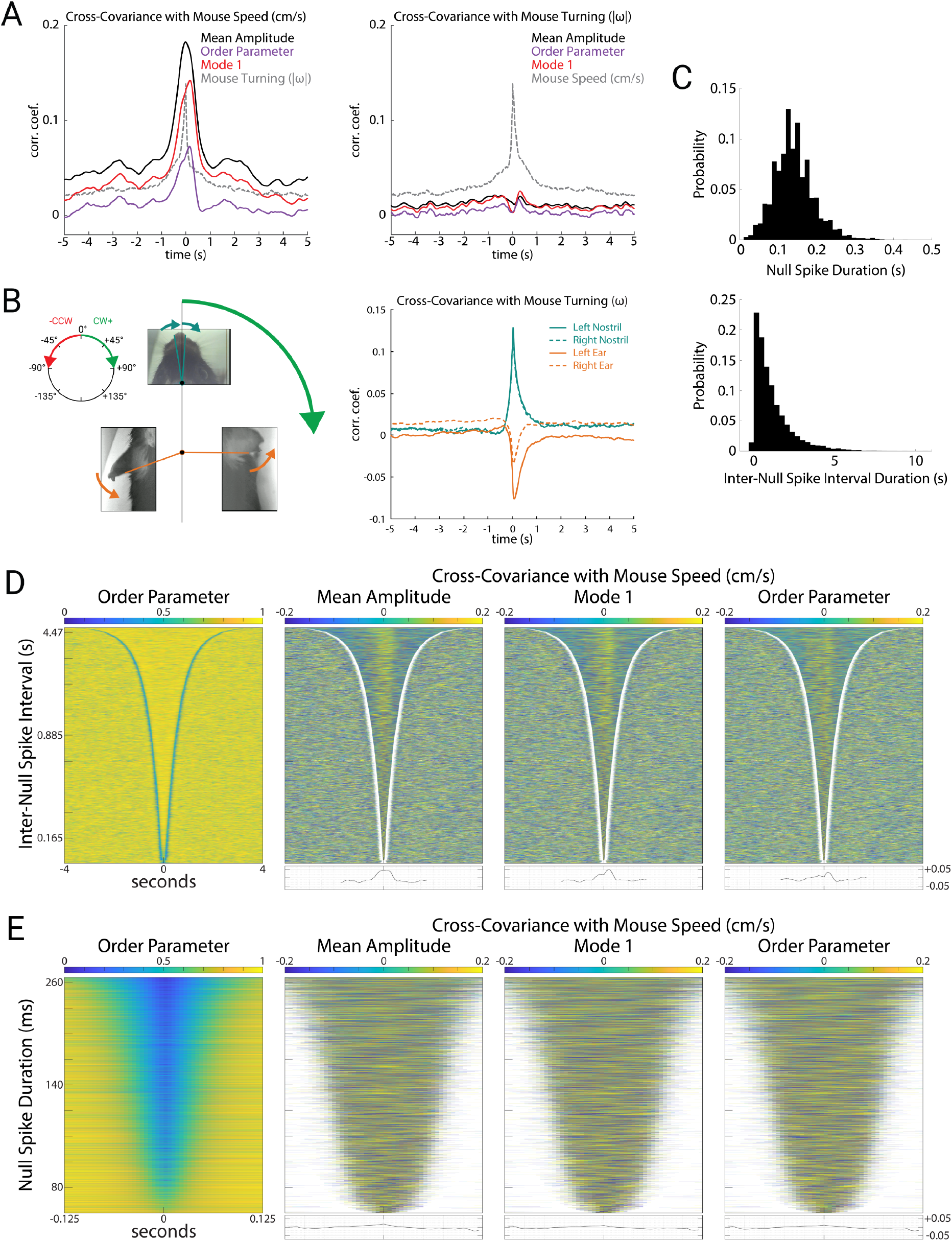
Mode 1 correlates with mouse speed during stable periods. A) Cross-covariance of the mean amplitude, order parameter, Mode 1 (percent variance explained), and mouse turning with mouse speed (left) and turning (right). Here we used the absolute value of angular velocity (|ω|, rads/sec) to avoid cancellation of correlations between clockwise and counterclockwise turns. B) Illustration of mouse nostril and ear movements (left). During a clockwise turn, we define mouse turning (ω) to be positive, and observed that motion of both nostrils strongly tended to also be clockwise (positive), whereas both ears strongly tended to move counterclockwise (negative). Cross-covariance of the nostrils and ears with mouse turning (right). Note the strong positive correlation of nostrils with turning, and the strong negative correlation of ear movements with turning (corresponding to the arrows depicted for the body parts at left). C) Distributions of null spike (top) and inter-null spike durations (bottom). D) Order parameter for all stable periods (N = 9212) between null spikes sorted by duration (left). Cross-covariance with mouse speed during each stable period for mean amplitude, Mode 1 (percent variance explained), and order parameter, with average cross-covariance of all stable periods plotted below (right). E) Order parameter for all null spikes (N = 9212) sorted by duration (left). Cross-covariance with mouse speed during each null spike for mean amplitude, Mode 1, and order parameter, with average cross-covariance of all null spikes plotted below (right)

We next wondered whether the prominent fluctuations in amplitude, Mode 1, and order parameter during null spikes underlie their correlation with mouse speed. To address this question, we detected null spikes using an algorithm based on the rate of change of the order parameter (see Methods). Null spikes (N = 9212) had a mean duration of 0.140 ± 0.051 s, and the stable periods between null spikes had a mean duration of 1.156 ± 1.164 s (Figure 4C), consistent with previous reports of null spikes in theta activity recorded from local brain regions (Kozma and Freeman, 2008). Figure 4D (left) shows the stable periods between null spikes, sorted by duration, such that the delineation of stable periods by the sharp drop in order parameter (null spikes) appears as a blue V-shaped boundary. Figure 4D (right) shows the covariance of mouse speed with three measures of theta activity (amplitude, Mode 1, and order parameter) during these stable periods. For all three measures, a positive correlation during the stable periods appears as a yellow band, and as a peak in the averaged covariance at the bottom. In contrast, mouse speed was not correlated with these theta measures during the null spikes themselves. Figure 4E shows all null spikes, sorted by duration. Covariance of mouse speed with amplitude, Mode 1, and order parameter were negligible during null spikes. This indicates that despite the large and coordinated fluctuations of amplitude, Mode 1, and order parameter during null spikes, these contribute little to the correlation between mouse speed and theta activity, which is instead driven by smaller fluctuations during the stable periods between null spikes.

### Null spikes impose a phase reset of Mode 1

Cortical null spikes thus appear to be independent of the correlation between theta oscillations and behavior. We therefore wondered what role they might play in the ongoing dynamics of the spatiotemporal modes of theta oscillations. The alternation of null spikes and the stable periods between them have been previously described as “beats” (Kozma and Freeman, 2008), suggesting that they could be a form of amplitude modulation arising from destructive interference between oscillations of slightly different frequencies.

To test this hypothesis, we examined phase slip across null spikes and stable periods. If amplitude modulation arises from beating between similar oscillation frequencies, the phase of the oscillation should be relatively stable and unaffected by the beats. Figure 5A shows a representative 5 second epoch with order parameter plotted at the top, and Modes 1 and 2 plotted underneath. Null spikes in the order parameter are color-coded in black, while the stable periods are color-coded by the phase of Mode 1 relative to an arbitrary reference sinusoid. Mode 1 phase was relatively constant during stable periods, but jumped sharply by up to 180° from one stable period to the next, across the intervening null spike. To visualize these Mode 1 phase jumps directly, we generated a “carry on” sinusoid for illustration purposes (thin black sinusoid in Figure 5A), using the phase angle and instantaneous frequency of Mode 1 just prior to the onset of the null spike, which then “carried on’’ for the null spike duration to serve as a reference for visualizing phase slip. This revealed that when Mode 1 reappeared following a null spike, it was typically out of phase from the time period just prior to the null spike.

**Figure 5:**
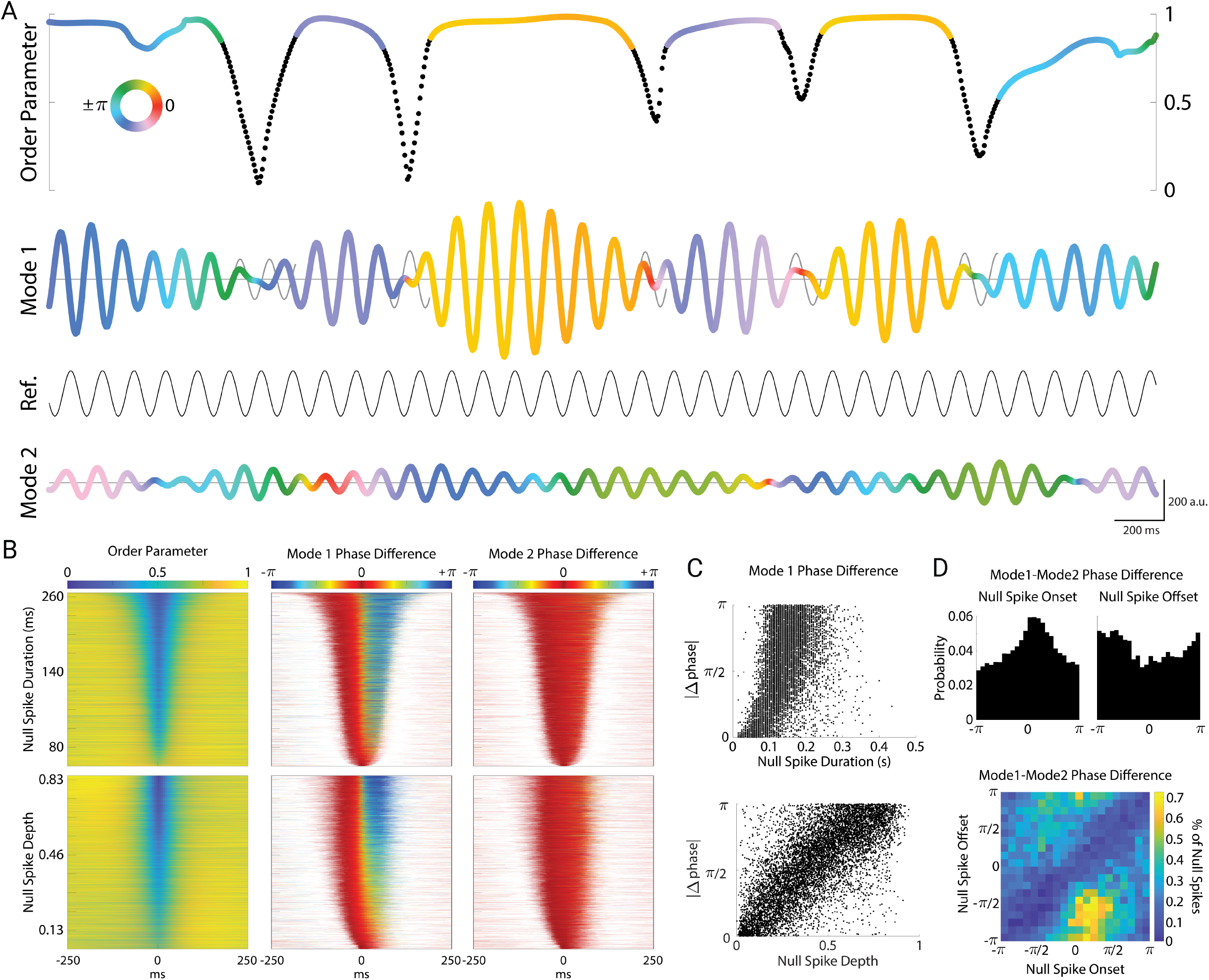
Cortical null spikes impose a phase-reset of Mode 1. A) Example plot of null spikes and corresponding phase resets of Mode 1. Color of the order parameter and Mode 1 correspond to the phase angle between Mode 1 and a reference sinusoid set at the average frequency of Mode 1 (8.7 hz). Black regions of the order parameter correspond to periods of a detected null spike. For each null spike, a “carry-on” signal is generated (gray sinusoids), representing the theoretical continuation of the mode at its instantaneous frequency immediately preceding the null spike. The color of Mode 2 corresponds to its own phase angle with the reference sinusoid. For simplicity, only the real components of the complex-valued modes are plotted (a.u.). B) All null spikes (N = 9212) sorted by duration (top row) and depth (bottom row). Corresponding phase differences of Mode 1 (middle) and Mode 2 (right) with their respective carry-on signals plotted for each null spike. C) Absolute value of change in phase plotted by null spike duration (top) and depth (bottom). D) Distribution of phase differences between Mode 1 and Mode 2 at the onsets (top left) and offsets (top right) of the null spikes. Density plot of the onset angles with their corresponding offset angles (bottom).

In contrast, the phase of Mode 2 (Figure 5A, bottom) varied smoothly and continuously and was unaffected by null spikes or stable periods. This was true across all null spikes and stable periods (Figure 5B). We sorted all null spikes by their duration (Figure 5B top left) to examine the phase slip of Mode 1 relative to the phase just prior to the onset of the null spike. The phase of Mode 1 jumped sharply at the midpoint of the null spike (Figure 5B top middle), whereas the phase of Mode 2 remained stable across the duration of the null spike (Figure 5B top right). When we instead sorted null spikes by their depth (peak-to-trough, Figure 5B bottom left), we observed that the magnitude of the Mode 1 phase jump during the null spike depended strongly on the depth of the null spike (Figure 5B bottom middle). That is, the deeper the null spike, the greater the phase jump across that null spike. Figure 5C shows the magnitude of the phase jump as a function of null spike duration (top) and null spike depth (bottom). While the magnitude of the phase reset showed a significant dependence on null spike duration (linear regression, F = 2271, p = 0, N = 9212 null spikes), an even stronger relationship was found with null spike depth (F = 10174, p = 0). Thus, the magnitude of phase reset was linearly related to the degree of cortical asynchrony, as well as the amount of time spent in the vortex-like state of the null spike.

Interestingly, null spikes tended to occur when Mode 1 and Mode 2 were nearly synchronized in phase. While null spikes occurred at various phase relationships, there was a higher probability of null spike onset at a phase difference near zero (with a peak at π/4 radians, Rayleigh’s z = 178, p = 10^−70^), and a corresponding trough at null spike offset (Figure 5D top). Because Mode 1 resets across null spikes while Mode 2 remains stable, null spikes effectively reset the phase relationship between Mode 1 and Mode 2. This is seen in the joint probability distribution of these two measures (Figure 5D bottom), which displays a prominent null-zone (blue diagonal), reflecting the fact that the phase relation is unlikely to remain the same at the onset and offset of the null spike. Null spikes occurring around the peak of π/4 radians typically resulted in phase reset to -2.2 radians (roughly 3π/4 radians). Thus, although only Mode 1 exhibits phase resets across null spikes, the two modes tend to have a particular phase relationship at the onset of null spikes.

## Discussion

We found that cortical theta oscillations are organized into distinct bilaterally-symmetric cortex-wide traveling waves. The dominant spatiotemporal pattern (Mode 1) displayed a divergent phase gradient originating in posterior retrosplenial cortex, radiating rostrolaterally across the cortex in a bilaterally symmetric manner. Secondary modes of oscillation (Modes 2 and 3) were bilaterally-symmetric spiral waves centered over somatosensory cortex. The rotational directions of Modes 2 and 3 were mirror-symmetric, and Mode 2 had high amplitude over medial frontal cortex whereas Mode 3 had high amplitude over the temporal poles. Mouse running speed was strongly correlated with mean theta amplitude, synchrony, and Mode 1 power. In contrast, other aspects of mouse behavior such as turning and movements of the ears and nose were unrelated to theta oscillations, even though these movements were correlated both with running speed and with each other. Theta oscillations showed metastable dynamics, in which stable periods of Mode 1 were punctuated by transient null spikes of low-amplitude asynchronous activity, during which Modes 2 and 3 were prominent. Moreover, these null spikes were associated with a phase reset of Mode 1. These brain-wide spatiotemporal patterns of theta oscillations could be involved in integrating sensory and motor information during natural behavior.

### Theta oscillations and traveling waves

In primates, theta oscillations coordinate large-scale cortical excitability, nested gamma oscillations, and mediate the rhythmic sampling of sustained attention (Bahramisharif et al., 2013; Helfrich et al., 2018; Lakatos et al., 2009). In rodents, theta-modulated spiking across the cortex has similarly established theta as a temporal organizer of activity (Fournier et al., 2020; Hafting et al., 2008; Holsheimer, 1982; Hyman et al., 2005; Muir and Bilkey, 1998; Siapas et al., 2005; Sirota et al., 2008). In contrast to synchrony, traveling waves have been hypothesized to allow some portion of the population to be responsive or optimally sensitive at any given time, and utilize distinct phase information to segment and categorize multiple inputs from each other and segment inputs from background (Ermentrout and Kleinfeld, 2001; Muller et al., 2018).

The spatial phase gradient of Mode 1 appears to resemble several patterns of neocortical organization observed across mammals. Mode 1 flows directly along the developmental expression gradients of Emx2 and Pax6, two highly conserved mammalian genes which regulate the arealization of the neocortex (Bishop et al., 2000). Additionally, this organization matches the principal gradient of functional connectivity observed in mice, which in turn reflects the evolutionary origins of the neocortex (Huntenburg et al., 2021). Lastly, Mode 1 is consistent with population analysis of traveling wave direction across the human cortex (Zhang et al., 2018), and congruent with the dorsal and ventral streams of feed-forward processing (Goodale and Milner, 1992). More recent evidence shows that the directions of cortical traveling waves change dynamically across behavior, reflecting distinct states of neural processing (Mohan et al., 2024).

### Metastability and phase resets

We found that theta oscillations occurred in bouts that lasted about 1 s, which were punctuated by brief ∼0.1 s periods of low amplitude and low synchrony, which we refer to as null spikes based on their similarity to previously described null spikes in gamma activity (Freeman, 2007; Kozma and Freeman, 2017). These findings are consistent with a growing number of reports of metastable dynamics, in which brain activity occurs as a sequence of discrete quasi-stationary states. Experimental and theoretical work has suggested that metastable states may underlie neural representations, with neural computation proceeding along a sequence of state transitions (La Camera et al., 2019; Recanatesi et al., 2022). Because our free exploration paradigm did not include an explicit task, we cannot relate the spatiotemporal motifs we observed to memory or perceptual representations. Nevertheless, our results suggest that metastable states may take the form of cortex-wide traveling waves, and that asynchronous spiral waves may underlie the rapid transitions between these metastable states. Moreover, these transitions appear to involve a phase reset in the dominant mode from one state to the next.

Phase resetting of ongoing oscillations modulates cortical excitability (Lakatos et al., 2009), and is critical for coding strategies that rely on phase information (Canavier, 2015; Voloh and Womelsdorf, 2016). Spontaneous cortical waves have been shown to be phase-reset through visual stimuli (Aggarwal et al., 2024), and phase resets of hippocampal theta oscillations have been observed to occur during critical moments of behavioral sequences such as jumping in rodents (Green et al., 2022), saccades in primates (Jutras et al., 2013), and working memory tasks in rats and humans (Givens, 1996; Tesche and Karhu, 2000). Here, we found that the magnitude of cortical null spikes was linearly related to phase resets of Mode 1. In other words, the amount of cortex-wide asynchrony during null spikes had a linear relationship to the resulting phase shift of the dominant cortical mode.

During null spikes, the spiral waves of Mode 2 remained stable, suggesting they are generated independently from Mode 1. Spiral waves are unique in that they necessarily contain an equal distribution of all phases at any given time, and therefore necessarily impose asynchrony. Therefore, as Mode 1 transiently weakened and order parameter dropped during a null spike, cortical theta became dominated by the asynchronous spiral waves of Modes 2 and 3.

The coupling of theta oscillations in frontal cortex with the cortex-wide spiral waves of Mode 2 may therefore reflect an important mechanism for bidirectional communication between the hippocampus and neocortex (Benchenane et al., 2010; Hyman et al., 2005; Jones and Wilson, 2005a, 2005b; Park et al., 2021; Siapas et al., 2005). Regardless of whether Mode 1 is coincident with, driven by, or a reflection of hippocampal theta (discussed in further detail below), phase-resetting in the presence of stable spiral waves across the cortex suggests the presence of at least one secondary global process. These spirals may reflect respiration-coupled oscillations, which have been observed to similarly entrain neurons across the cortex, and particularly through the frontal cortex (Karalis and Sirota, 2022; Tort et al., 2018).

Alternatively, previous studies have theorized that spiral waves may provide a means for the cortex to flexibly produce emergent ongoing oscillations in the absence of an explicit pacemaker, as well as allow populations to disengage from globally-synchronous rhythms and produce a state change (Huang et al., 2010). These potential functions appear consistent with our findings of ongoing cortex-wide spiral waves and their stability across the rapid state transitions during null spikes. The large-scale spiral waves seen here may therefore provide the means for multimodal processes to emerge and assimilate into ongoing activity across the neocortex in a spatiotemporal manner. Spiral waves would specifically serve as an ideal scaffold for large-scale phase coding, due to their unique ability to enforce exactly one cycle (2π) across neural space, thereby maximizing phase separability and preventing phase redundancy (Ermentrout and Kleinfeld, 2001). Such a property would most likely be vital to integrate many disparate stimuli and ongoing processes into a single flexible cortical framework.

### Possible sources of cortical theta

There are at least 3 mechanisms that could generate the spatiotemporal patterns of theta oscillations in cortex, which are not mutually exclusive.

First, they could be produced by volume conduction from a hippocampal theta generator. This possibility is consistent with the spatial amplitude pattern of Mode 1, which is greatest over parietal cortex, where the dorsal aspect of CA1 is closest to the overlying cortex. The dependence of theta amplitude and synchrony on mouse speed is also consistent with a theta generator in the hippocampus, where theta shows a similar dependence on speed. Indeed, a previous report of widespread cortical theta oscillations concluded that it most likely arises by volume conduction from the hippocampus (Sirota et al., 2008); see also (Gerbrandt et al., 1978; Kennedy et al., 2023). However, subsequent studies revealed that hippocampal theta oscillations are organized into a traveling wave along the septotemporal axis (Lubenov and Siapas, 2009; Patel et al., 2012; Zhang and Jacobs, 2015). This direction of travel is orthogonal to the rostrolateral propagation of the Mode 1 traveling wave, which is therefore inconsistent with the possibility that Mode 1 is explained purely by volume conduction. The spatial amplitude patterns and spiral wave structures of Modes 2 and 3 are also inconsistent with volume conduction from the hippocampus, as are their lack of dependence on mouse speed.

A second possible mechanism that could underlie cortical theta oscillations is that the rhythm originates in the hippocampus, but that field potential oscillations are generated in cortex by synaptic input arising from widespread projections from the hippocampus to neocortex. In this scenario, cortical theta is hippocampally driven, but could exhibit a different spatiotemporal pattern due to transformation by hippocampal axonal projection patterns. This could explain the different directions of traveling waves in hippocampus and cortex, but cannot explain the existence of the additional spatiotemporal patterns exhibited by Modes 2 and 3.

Finally, a third possible mechanism that could underlie cortical theta oscillations is local generation and propagation by intracortical circuitry, either as an emergent cortical process or by entrainment from hippocampal input (Zold and Hussain Shuler, 2015). Because we did not have a depth electrode in the hippocampus, we cannot directly relate cortical theta oscillations to hippocampal theta. However, parietal theta (which we found to have the highest amplitude in Mode 1) has been consistently shown to be strongly coherent with that of the hippocampus (Gerbrandt et al., 1978; Sirota et al., 2008; Tort et al., 2018), providing further evidence that Mode 1 is strongly related to hippocampal theta.

In contrast, the identity of Modes 2 and 3 as a distinct extrahippocampal process are supported by their lack of coherence with Mode 1, stability during phase-resets, and their distinct amplitude maps and spiral structure. Spiral waves have been described in humans during sleep spindles (Muller et al., 2016). In mice, brain-wide spiral waves in the 2-8 Hz frequency range (below our 9 Hz theta frequency) have also been recently described (Ye et al., 2023). These spirals are centered over somatosensory cortex, in agreement with our observations (Fig. 2D).

Interestingly, that study reported a circular arrangement of axonal projections in sensory cortex, which matched the spiral waves we and they observed, providing a possible anatomical basis for the structure and location of these spirals over somatosensory cortex. This provides further evidence that these spiral waves (Modes 2 and 3) may be generated and propagated by intracortical circuitry rather than being driven by hippocampal theta.

Future experiments pairing single unit recordings with large-scale electrocorticography will allow the investigation of how spiking activity relates to these global oscillatory modes, and distinguish between the possible mechanisms described above. The large-scale theta oscillations we describe here could be involved in the dynamic routing of information across widespread cortical areas. The integration of sensory and motor information during ongoing natural behavior requires a system for flexible communication among disparate cortical areas. Dynamic and metastable sequences of spatiotemporal theta motifs could provide a substrate for such flexible communication.

## Figure legends

Supplementary Video 1: Cortical cyclograms of the top three modes, showing the progression of phase along their spatial phase gradients across three full theta cycles (top and bottom rows). The red line in the middle row indicates the current phase angle.

## Methods

All procedures were in accordance with the National Institutes of Health guidelines, as approved by the University of Oregon Animal Care and Use Committee.

### Electrode arrays and headset

Epidural electrode arrays were composed of stainless steel 000-120 flathead slotted drive screws (Antrin Miniature Specialties) that we soldered to 42 mm long teflon-coated stainless steel wire (A&M systems, recording electrodes: 790500 50.8μm diameter, ground and reference electrodes: 791400 127μm diameter) and pinned to a 32-channel electrode interface board (Neuralynx 31-0603-0127). Reference and ground channels were soldered to one common screw. Impedances were confirmed to be within the range of 1-3Ω for the ground and reference channels, and 16-17Ω for the recording channels. The spatial layout of the electrode array was adapted from the epicranial design of (Mégevand et al., 2008), with the inter-electrode distance increased to 1.8 mm, to fully span the cortex when curved over its surface. A stencil of electrode locations was laser-cut from filter paper (Whatman 1001-185), for use during surgery. The carrier headset for head-mounted camera arrays (Sattler and Wehr, 2020) was mounted to a 32-channel headstage (Intan #C3324).

### Surgical procedure

Mice were anesthetized with isoflurane (1.5%). The electrode array stencil was aligned to bregma, flattened over the skull, and used to mark electrode locations on the surface of the skull. Craniotomies were then created over each electrode location to fully expose the dural surface, and the electrode was fastened into place and covered with dental acrylic. Mice were housed individually after surgery, and allowed at least a week of postoperative recovery before recordings.

### Animals and experimental procedure

We recorded from C57Bl6 mice (N = 3 mice, 2 male and 1 female). Experiments were performed in an electrically shielded sound-attenuating chamber. Mice were placed in a 60 cm diameter cylindrical arena for 2 exploration sessions a day (1 in the dark (5 minutes) and 1 in the light (5 minutes)) for 6 days. Mice were food restricted for the last three days (1 hour of access to food per day, after their sessions). Mice were active for all sessions (locomoting or rearing), with rare periods of grooming or behavioral quiescence. Sessions were halted if the tethers of the mouse became tangled, and were resumed for the remaining time in each session once the tethers had been untangled.

### Data acquisition

Continuous data was recorded at 30 kHz sampling rate using a 32-channel headstage (Intan Technologies #C3324), USB interface board (Intan Technologies RHD2000 board), and Open Ephys software (Siegle et al., 2017) with a digital bandpass filter of 2.4959 to 7603 Hz.

The overhead camera (FLIR BFS-U3-16S2M) was captured at 200 Hz, and the head mounted cameras were captured and deinterlaced at their 60 Hz field-rate using Bonsai (bonsai-rx.org). Signals were synchronized using illumination of an IR LED from TTL pulses as described in (Sattler and Wehr, 2020).

### Behavioral tracking

All behavioral data was tracked using DeepLabCut (Mathis et al., 2018). From the overhead camera, six points on the head-mounted camera headset of the mice were used to calculate their bearing in the arena, and a point on the back of their neck was used to calculate their speed. From the mounted head camera, the left and right nostrils were tracked, and their bearing relative to the mouse’s midline were calculated. From the ear cameras, a distinct point on the posterior edge of the pinna was tracked, and their bearing relative to the mouse’s midline were similarly calculated.

### Pre-analysis

For all analysis, the raw continuous data was first bandpass filtered at 1 to 30 Hz using a 1st order butterworth filter (implemented using *filtfilt*.*m* in Matlab), and downsampled to 200 Hz to facilitate data processing while retaining high fidelity. All behavioral measurements from the cameras were similarly resampled at 200Hz. For two mice, a dead channel was interpolated from neighboring channels prior to downsampling. Sessions which contained an instance of a large electrode artifact (4 sessions from 1 mouse) were excluded from analysis.

### Analytic signals

Data was filtered at 9 Hz using wavelet-based filtering (implemented using *wFilter*.*m* from Matlab Central File Exchange), with a bandwidth of 1 standard deviation of the wavelet’s gaussian function, and converted into analytic signals using a Hilbert transformation.

Order parameter (r) was calculated as:

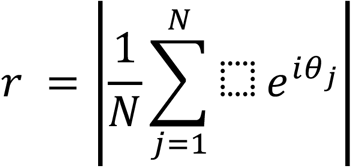

where N = number of oscillators (i.e., electrodes), and *θ* = phase. Null spikes were detected by identifying periods in which the absolute value of the change in order parameter exceeded 0.015/s.

### Singular value decomposition

Singular value decomposition (SVD) was performed on the complex-valued analytic signals extracted from the theta-filtered LFPs to produce eigenmodes of oscillatory activity as follows:

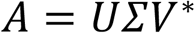

where *A* is an *n* by *t* matrix of *n* analytic signals at *t* timepoints, *U* is an *n* by *n* complex-valued matrix where each column represents the eigenvectors, *Σ* is an *n* by *t* diagonal matrix containing the singular values for each eigenvector, *V* is a *t* by *t* complex-valued matrix where each row represents the eigenvalues at each timepoint for each eigenmode, and * represents the Hermitian transposition.

Additionally, an arbitrary “reference” channel was chosen after decomposition to align each eigenvector (and associated eigenvalues) to a common phase angle. We chose the right posterior channel of retrosplenial cortex (inter-electrode coordinate: -4.2288 AP, +0.9 ML), and rotated each eigenvector (and their respective eigenvalues) as follows:

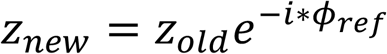

Where *z*_*old*_ is the original complex-valued eigenvector (or eigenvalue), *ϕ*_*ref*_ is the initial phase angle of the reference channel of the eigenvector, and *z*_*new*_ is the resulting complex-valued eigenvector (or eigenvalue). This therefore aligned the eigenmodes such that the chosen reference channel was at a phase angle of 0, and shifted their respective eigenvalues appropriately.

The amount of variance explained by each successive eigenmode decreased smoothly and continually for modes 2 and beyond, with no abrupt drop off in variance explained. Here we decided to limit our analysis to the first 3 modes, because the spatiotemporal patterns of the first 3 modes were consistent across mice, whereas the spatiotemporal patterns of modes 4 and beyond were variable across mice. We performed SVD on the combined data from all mice and analysis.

## Statistics

Pearson’s correlation coefficients (*r*) were computed separately for each behavioral trial, and then averaged across trials. For significance testing of correlation coefficients across all trials, we used the Wilcoxon signed-rank test to compare the correlation coefficient for each trial to a control correlation coefficient computed by time-reversing one of the signals. This permutation control preserves the autocorrelation structure of each signal but removes the temporal relationships between them.

To test the relationship between null spike onsets and Mode 1-2 phase coupling (Fig. 5D), we binned phase differences between Mode 1 and Mode 2 across the dataset in 25 bins from -π to π, and then normalized the null spike count in each bin to the background count of Mode 1-2 phase differences in that bin (across time). We then tested for the presence of a unimodal peak using the Rayleigh test for non-uniformity of circular data.

## Spatial phase gradient

Spatial phase gradients were calculated for each channel using its set of nearest neighbors, and therefore consisted of a scalar value (radians/mm) for each of the three spatial components of the hexagonal lattice: x (−π,0), y (−2π/3,π/3), and z (−π/3,2π/3). Phase derivatives were taken for each of these components by taking the angle of the product of the analytic signal with the complex conjugate of the second, using the forward or central difference when available. Vector summation of these three components was then used to calculate the resultant vector. To calculate curl, the spatial vector field was interpolated into a 15×15 matrix, and quantified using *curl*.*m* in Matlab.

## Data and Code Availability Statement

Data and code are available upon request.

## Author Contributions

NJS performed research and analyzed data. NJS and MW designed research and wrote the paper.

